# CHMP4B contributes to maintaining the follicular cells integrity in the panoistic ovary of the cockroach *Blattella germanica*

**DOI:** 10.1101/2024.01.24.577052

**Authors:** Farrus, J.L. Maestro, M.D. Piulachs

**Affiliations:** Institut de Biologia Evolutiva (CSIC-Universitat Pompeu Fabra), Passeig Maritim de la Barceloneta, 37. 08003 Barcelona, Spain

**Keywords:** Insect oogenesis, insect reproduction, shrub, snf7, Vps32, ESCRT-III, insect ovary

## Abstract

The Endosomal Sorting Complex Required for Transport (ESCRT) is a highly conserved cellular machinery essential for many cellular functions, including transmembrane protein sorting, endosomal trafficking and membrane scission. CHMP4B is a key component of ESCRT-III subcomplex and has been thoroughly studied in the meroistic ovaries of *Drosophila melanogaster* showing its relevance in maintaining this reproductive organ during the life of the fly. However, the role of the CHMP4B in the most basal panoistic ovaries remains elusive. Using RNAi, we examined the function of CHMP4B in the ovary of *Blattella germanica* in two different physiological stages: in last instar nymphs, with proliferative follicular cells, and in vitellogenic adults when follicular cells enter in polyploidy and endoreplication.

In *Chmp4b*-depleted specimens, the expression of actin in the basal pole of the follicular cells increased, leading to an excess of actin bundles that surrounded the basal ovarian follicle and modifying their shape. Depletion of *Chmp4b* also caused a loss of planar polarity in the follicular epithelium due to the actin increase in cell membranes, resulting in different cell morphologies and sizes, as well as an absence of patency. In these cells, the nuclei appeared unusually elongated, suggesting an incomplete karyokinesis.

These results proved CHMP4B essential in preserving the proper expression of cytoskeleton proteins vital for basal ovarian follicle growth and maturation, and for yolk proteins incorporation. Moreover, the correct distribution of actin fibers in the basal ovarian follicle emerged as a critical factor for the successful completion of ovulation and oviposition. The overall of the results, obtained in two different proliferative stages, suggest that the function of CHMP4B in *B. germanica* follicular epithelium is independent of the proliferative stage of the tissue.

## INTRODUCTION

The Endosomal Sorting Complex Required for Transport, also known as ESCRT, consists of five protein subcomplexes ( ESCRT-0, -I, -II, -III, and VPS4, see Christ et al., 2017) highly conserved between yeast, humans, and insects (Li & Blissard, 2015). ESCRT subcomplexes are required for transmembrane protein sorting, endosomal trafficking, membrane remodeling, and membrane scission (Christ et al., 2017). CHMP4B (charged multivesicular body protein 4B) from humans (Peck et al., 2004), the orthologue of Snf7/VPS32 in yeast (Kranz et al., 2001) and of Shrub (Shrb) in *Drosophila melanogaster* (Sweeney et al., 2006), is the main component of ESCRT-III subcomplex. CHMP4B plays a crucial role in membrane protein trafficking and executes membrane scission during processes such as multivesicular body formation and cytokinesis, helping to separate the daughter cells (Carlton & Martin-Serrano, 2007; Christ et al., 2017).

In *Drosophila melanogaster* ovaries, ESCRT complexes are not essential for the global oocyte development (Vaccari et al., 2009). They are involved in the endosomal arrangement of membrane proteins, including the localization of Notch receptor, which signaling is essential for the development of the follicular cells and oocyte maturation (Klusza & Deng, 2010). Also, Shrb participates in the membrane abscission during cytokinesis of germinal stem cells, where it is recruited at the midbody, the thin intercellular membrane bridge that connects the two daughter cells. The interaction between Shrb and ALiX proteins ensures cytokinesis completion (Christ et al., 2017; Eikenes et al., 2015; Matias et al., 2015; Raiborg & Stenmark, 2009). Moreover, *D. melanogaster ESCRT* mutants lack the subcortical actin in germinal cells. In general, mutants for any of the ESCRT components show strong defects in actin and plasma membrane organization that can result in multinuclear cells (Vaccari et al., 2009).

These actin flaws have led to the proposal that ESCRT complexes could directly regulate the actin cytoskeleton (Sevrioukov et al., 2005; Vaccari et al., 2009). The Shrb function was also analyzed in *D. melanogaster* males. It was observed that depletion of *shrb* mRNA in fly testes determines a significant increase in the expression of certain cytoskeleton-associated genes. Results pointed to this increase as the responsible factor for the arrest of germ cell development, which led to the accumulation of early gametes that did not reach maturation (Chen et al., 2021).

The study of the function of Shrb in insect ovaries is limited to *D. melanogaster*, a species with a meroistic polytrophic ovary type, where the nurse cells (from germinal lineage) escort the oocytes during the egg chamber development. In the present work, we study the function of Shrb/CHMP4B in a basal ovary type, the panoistic ovary, using as an experimental model the cockroach *Blattella germanica*. The panoistic ovary lacks nurse cells and the oocyte is responsible for synthesizing the RNAs and proteins that will be necessary to determine future embryo development. After leaving the germarium, early differentiated oocytes are surrounded by a monolayer of somatic follicular cells (FC), forming an ovarian follicle (Rumbo et al., 2023). In *B. germanica* oogenesis, only the basal oocyte in each ovariole matures in a gonadotropic cycle. The other ovarian follicles in the ovariole remain arrested in the vitellarium, waiting for the next cycle (Irles & Piulachs, 2014). The basal ovarian follicles (BOFs) become active in the sixth (last) nymphal instar when the oocyte is synthesizing RNAs and proteins that will be necessary for the oocyte to mature and for future embryo development (Elshaer & Piulachs, 2015; Irles & Piulachs, 2014). At the same time, the follicular epithelium in the BOF proliferates actively, producing an adequate number of cells distributed homogeneously to cover the oocyte surface. In 3-day-old adults, coinciding with the increase of juvenile hormone in the hemolymph and the beginning of the vitellogenic period, the follicular epithelium in the BOF reaches the maximum number of cells, changing their program and activating the mitosis-endocycle switch (Irles et al., 2016). In 5-day-old adults, all the FCs are binucleated, become polyploidy, and enter into endoreplication (Irles & Piulachs, 2014; Zhang & Kunkel, 1992). Then, the chorion synthesis starts.

During vitellogenesis, the oocytes in the BOFs take up proteins from the hemolymph to grow exponentially (Bellés et al., 1987). To allow these storage proteins to reach the oocyte membrane, the FCs contract their cytoplasm, leaving large intercellular spaces between them in a process called patency. This process, under juvenile hormone control, was first described in *Rhodnius prolixus* (Davey, 1981; Davey & Huebner, 1974). Recently, the mechanism behind this process was studied in *D. melanogaster* (Isasti-Sanchez et al., 2021; Riechmann, 2021). According to these studies, the release of tension in the actin-myosin cytoskeleton and the removal of the adhesion proteins that participate in cell junctions determine a change in the number of connections between the FCs, which pass from tricellular junctions to bicellular junctions, leaving spaces between cells (Isasti-Sanchez et al., 2021). At the end of vitellogenesis, the FCs close these spaces, and the gonadotropic cycle ends with the chorion synthesis just before ovulation and oviposition.

*B. germanica* allow us to study the function of Shrb/CHMP4B on the FCs in different situations: in the last instar nymphs, when FCs are actively proliferating, increasing their number as the BOF grows, and in adults, when the proliferative stage of FCs finishes, the cells become binucleated and enter into polyploidy and endoreplication. We found that Shrb/CHMP4B in the panoistic ovary of *B. germanica* regulates the actin cytoskeleton and associated genes involved in maintaining the follicular epithelium planar polarity. It is necessary to maintain the right levels of Shrb/CHMP4B in the ovary to correctly execute the changes of the FCs program in the BOF and thus ensure their correct growth.

## MATERIAL AND METHODS

### Insects sampling

Specimens of the cockroach *B. germanica* were obtained from a colony fed on dog chow and water *ad libitum*, kept in the dark at 29 ± 1 °C and 60 – 70 % relative humidity. Dissections and ovarian tissue sampling were performed on carbon dioxide-anaesthetized specimens, held under Ringer’s saline. After dissection, tissues were immediately frozen in liquid nitrogen and stored at -80 °C.

### RNAi experiments

To deplete the expression of *shrb*/*Chmp4b* (from now *Chmp4b*), a dsRNA (ds *Chmp4b*) was designed targeting the conserved domain of the Snf7 superfamily (461bp). A heterologous 441 bp fragment from the gene sequence of the *Autographa californica* nucleopolyhedrovirus was used as negative control (dsMock) (Lozano & Belles, 2011). The dsRNA was synthesized using MEGAscript™ RNAi kit (Invitrogen), following the manufacturer instructions and stored at −20 °C until its use. The dsRNA was injected at a dose of 1 μg (1 μg/μL) into the abdomen of freshly emerged last (sixth)-instar female nymphs (N6D0) and newly emerged adult females, using a Hamilton microsyringe (Teknokroma). Sixth nymphal instar ovaries were dissected at day six of the instar. Ovaries from adult individuals were dissected on day 0 or 5 of the first gonadotrophic cycle, depending on the experiment.

### RNA extraction, cDNA synthesis and quantitative real-time PCR analysis

Total RNA from the ovaries was extracted using HigherPurity™ Tissue Total RNA Purification kit (Canvax Biotech S.L.). A total of 500 ng of RNA were then retrotranscribed using the Transcriptor First Strand cDNA Synthesis kit (Roche LifeScience) as previously described (Montañés et al., 2021). The absence of genomic contamination was confirmed using a control without reverse transcription.

The expression levels of the different genes studied were analyzed by quantitative real-time PCR (qRT-PCR) using cDNA from ovaries. The schedule used for the amplifying reaction was as follows: (i) 95 °C for 3 min, (ii) 95 °C for 10 s; (iii) 57 °C for 1 min; (iv) steps (ii) and (iii) were repeated for 445cycles. The expression of *actin-5c*, *armadillo* ( *arm*), *caspase-1* ( *casp-1*), *Chmp4b*, *dachsous* (*ds*), *eukaryotic translation initiation factor 4aIII* (e *IF4aIII*), *extracellular serine/threonine protein kinase four-jointed* (*fj*), *frizzled* (*fz*), *kugelei* (*kug; fat2* in *D. melanogaster*), *Myosin heavy chain* (*Mhc*) and protocadherin-like wing polarity protein *starry night* (*stan*) were quantified from 3-6 independent biological samples, making three technical replicates of each one, in a 10 μL of final volume. *Actin-5c* was used as a reference gene to obtain the expression profile of *Chmp4b* in ovaries, and e*IF4aIII* was used as a reference gene to compare treated and control samples. The sequence of the primers used, and the accession number of genes analyzed are detailed in Table S1.

### Tissue staining and immunohistochemistry

Ovaries from 6-day-old last instar nymphs and from 0-day- and 5-day-old adults were dissected and immediately fixed in paraformaldehyde (4% in PBS) for 2 h. Washing samples and antibody incubations were performed as previously described (Irles & Piulachs, 2014). The primary antibody employed was rabbit anti-Shrb (a gift from Dr. Thomas Klein; Bäumers et al., 2019) was used at dilution 1:50. The secondary antibody used was Alexa-Fluor 488 goat anti-rabbit IgG (Molecular Probes). In addition, ovaries were incubated at room temperature for 20 min in 300 ng/mL phalloidin-TRICT (Sigma) for F-actin staining and then, for 5 min in 1 µg/mL of DAPI (Sigma) for DNA staining. Ovaries were mounted in Mowiol (Calbiochem) and observed using a Zeiss AxioImager Z1 microscope (Apotome) (Carl Zeiss MicroImaging).

### Statistical analysis

All data were expressed as mean ± standard error of the mean (S.E.M.) from at least three independent experiments. The data were evaluated for normality and homogeneity of variance using the Shapiro–Wilk test, which showed that no transformations were needed. All datasets passed normality test. Statistical analyses were performed employing Student’s *t*-test. A p value < 0.05 was considered statistically significant. Statistical analyses were performed using GraphPad Prism version 8.1.0 for Windows, GraphPad Software.

## RESULTS

### *Chmp4b* expression in the ovaries of *Blattella germanica*

*Chmp4b* mRNA is highly expressed in ovaries during all the sixth nymphal instar, showing the highest values in the first day of the instar, just after the molt. Then, its expression in ovaries begins to decline reaching the lowest values in 5-day-old nymphs, coinciding with the highest levels of ecdysteroids in the hemolymph. Just before the imaginal molt *Chmp4b* mRNA levels were recovered coinciding with the decrease of ecdysteroids (Figure 1A). After the molt to adult, expression of *Chmp4b* mRNA in ovaries quickly decreased until 3-day-old females, when the vitellogenic period starts and FCs arrest cytokinesis to become binucleated. These relatively low levels of *Chmp4b* mRNA were maintained stable until the end of the gonadotropic cycle (Figure 1A).

**Figure 1.**
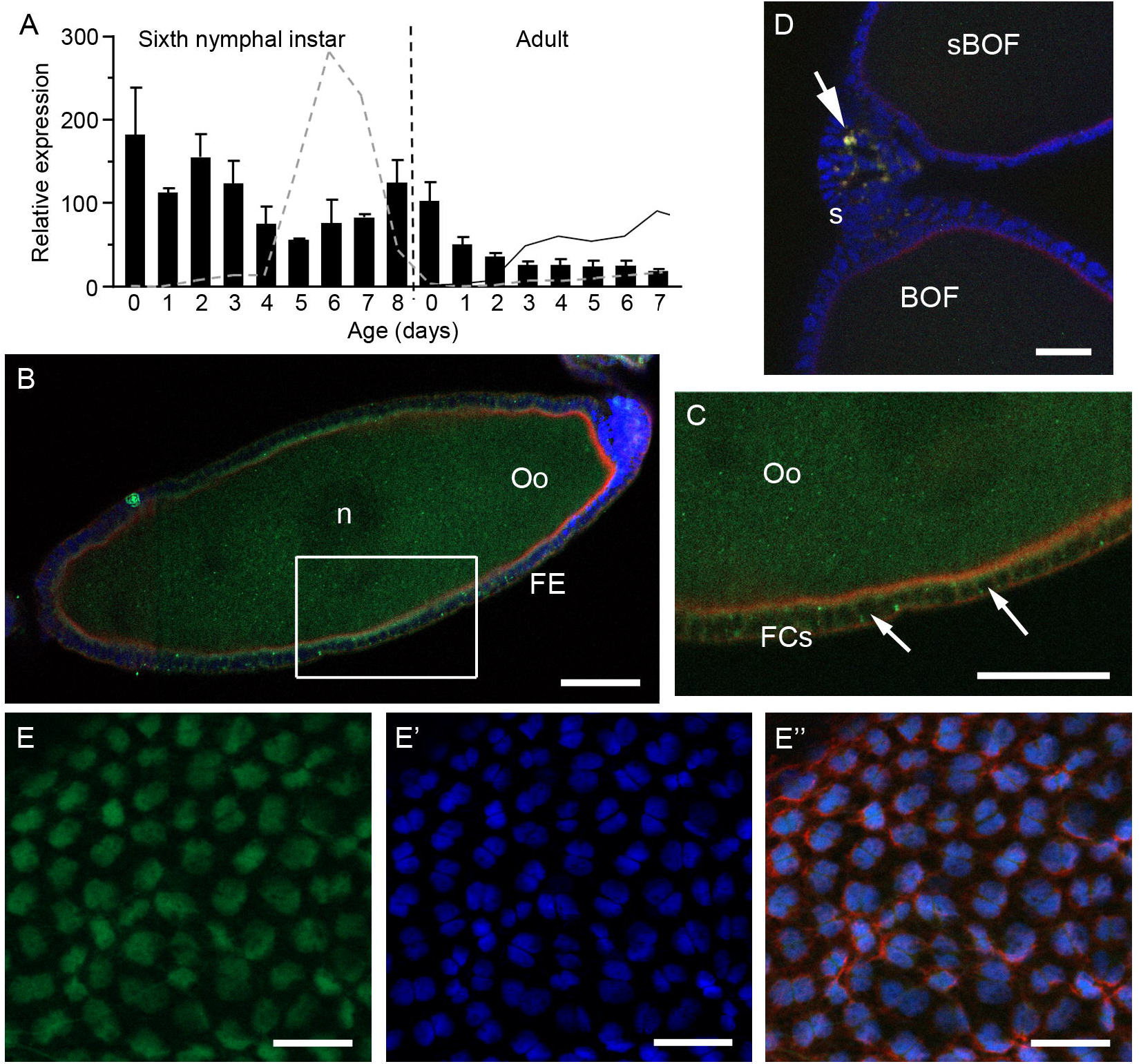
CHMP4B in *Blattella germanica* ovaries. (A) mRNA expression pattern of *Chmp4b* in ovaries of sixth nymphal instar and adults during the first reproductive cycle. The black dashed line indicates the molt to adult. Profiles of ecdysteroid titer in hemolymph (grey dashed line) and ecdysteroid contents in adult ovary (solid black line) are indicated (data from Cruz et al., 2003; Romaña et al., 1995). Data represent copies of *Chmp4b* mRNA per 1000 copies of *actin-5c* (relative expression), and is expressed as the mean ± S.E.M. (n = 3-4). (B) Immunolocalization of CHMP4B (green) in the BOF from a 0-day-old adult, showing the labelling spread by the oocyte cytoplasm. The oocyte apical pole is at the bottom-left of the image. (C) Detail of B showing the CHMP4B labelling in the cytoplasm of follicular cells and not in the nuclei (arrows). The images B and C were overexposed (500ms) to detect the fluorescence for CHMP4B in the corresponding channel. FE: Follicular epithelium; FCs: Follicular cells; n: nucleus; Oo: Oocyte. (D) Stalk (s) between the basal (BOF) and sub-basal ovarian follicles (sBOF) of 0-day-old adults, showing labeling for CHMP4B in the apical pole of the stalk cells (arrow). (E) Follicular epithelium in BOF from a 5-day-old adult showing CHMP4B localized in the nuclei of FCs. (E’) DAPI (blue) staining of nuclei from image E. (E’’) Merge image of E and E’ showing the nuclear localization of CHMP4B at this age. F-actin microfilaments were stained with phalloidin-TRITC (red). Scale bar in B and C: 50 µm, in D: 20 µm and in E: 50 µm.

Using a heterologous antibody against Shrb of *D. melanogaster* (Bäumers et al., 2019), we localized CHMP4B in ovaries of *B. germanica* adults. In newly emerged females, labelling for CHMP4B was very faint, and an over exposition was necessary to see the labelling in the cytoplasm of both, the basal oocyte and FCs from BOF (Figure 1B). The labelling, at this age, was absent from the FC nuclei (Figure 1C). Labelling for CHMP4B appeared also in the apical pole of the cells that are forming the stalk between the BOF and the sub-basal ovarian follicles (sBOF) (Figure 1D). Later, in 5-day-old adult females, labeling for CHMP4B in ovaries was localized in the nuclei of the FCs from the BOFs (Figure 1E-E’’), a translocation of the labeling that can be related with the change in the FCs program that occurs at this age. Due to the big oocyte size, we cannot observe the antibody labelling inside the basal oocyte of 5-day-old females.

### Effects of *Chmp4b* depletion on the ovary of 5-day-old *Blattella germanica* adults

*Chmp4b* functions related to oocyte development in *B. germanica* were studied using RNAi. Newly emerged adult females were treated with ds *Chmp4b* or dsMock, and the expression levels of *Chmp4b* were analyzed in ovaries from 5-day-old adults (n = 3, for each treatment). The *Chmp4b* expression was significantly depleted (p < 0.01, Figure 2A), reaching a 49% of reduction compared to dsMock. We measured *actin-5c* mRNA levels which were significantly upregulated (p < 0.01, Figure 2B), showing a two-fold increase compared to dsMock treated females indicating that also in *B. germanica* CHMP4B could regulate actin cytoskeleton expression.

**Figure 2.**
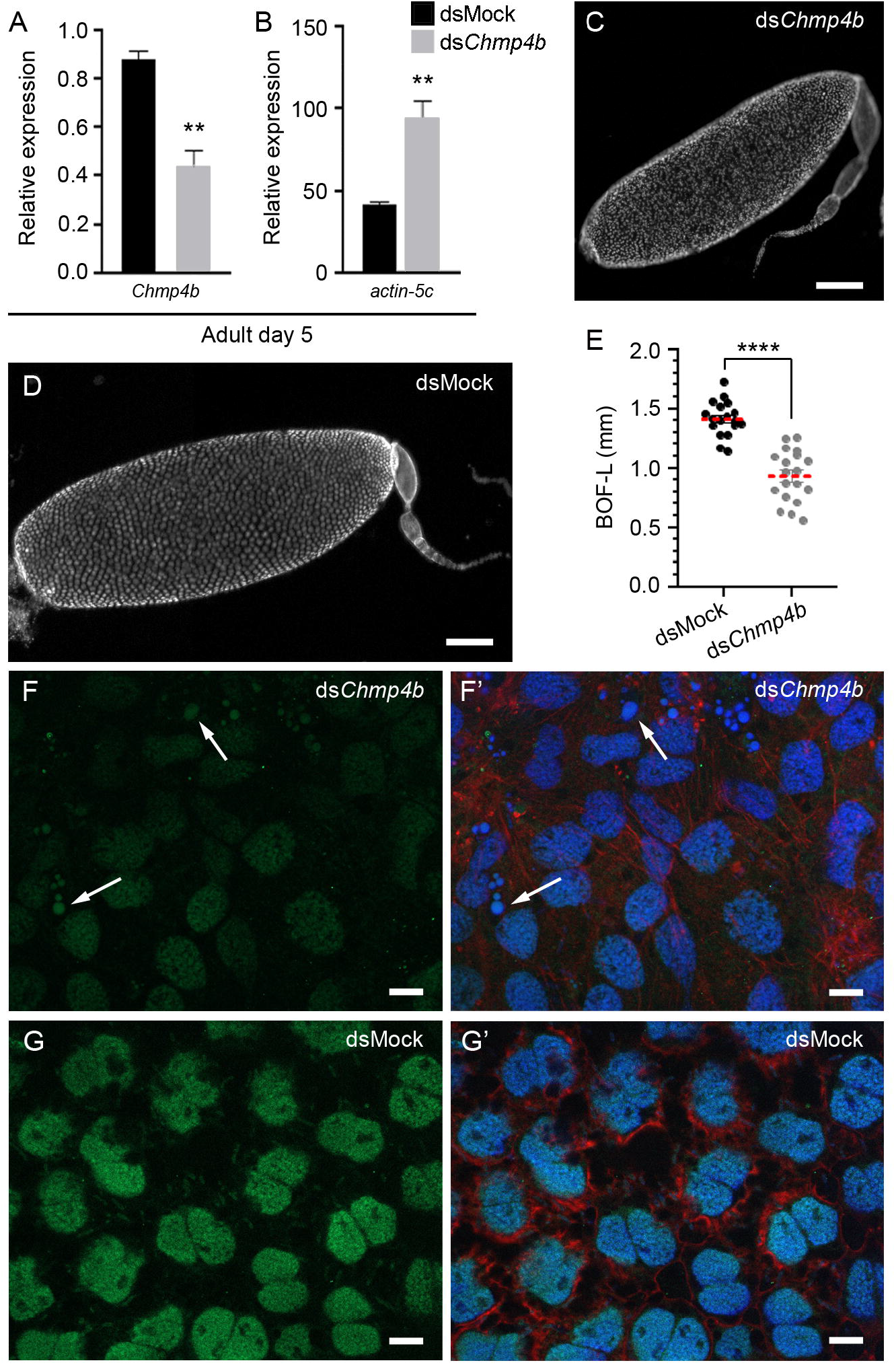
CHMP4B depletion in 5-day-old *Blattella germanica* ovaries. Newly emerged adult females were treated with ds *Chmp4b* or dsMock and dissected 5 days later. (A) Relative expression of *Chmp4b* in ovaries, showing the significant decrease of *Chmp4b* mRNA (n =3; ** p = 0.0047). (B) Relative expression of *actin-5c* in ovaries, showing the significant increase of *actin-5c* (n =3; ** p = 0.0093). Data represent copies of *Chmp4b* mRNA per copy of e*IF4aIII* mRNA (relative expression). Data is expressed as the mean ± S.E.M. (C) Ovariole from ds *Chmp4b*-treated adult. (D) Ovariole from dsMock-treated adult. In C and D, DNA was stained with DAPI (white). (E) Length of basal ovarian follicle (BOF-L; mm) in dsMock- and ds*Chmp4b*-treated females (n = 18 and 19 females, respectively; 3 – 5 ovarian follicles were measured per insect; **** p < 0.0001). The red dashed line indicates the mean. (F) Immunolocalization of CHMP4B in the nuclei of the follicular cells of ds *Chmp4b*-treated females (green). This image was overexposed until some signal was detected in the corresponding channel (500 ms of exposition). (F’) Merged image showing the CHMP4B labelling, and the F-actin and DNA staining. The arrows in both images indicate vesicles containing rests of DNA. (G) Immunolocalization of CHMP4B in follicular cells of dsMock-treated females (320 ms of exposition). The labelling is limited to the nuclei of follicular cells (green). (G’) Merged image showing the CHMP4B labeling, and the F-actin and DNA staining. CHMP4B was detected using a rabbit anti-Shrb antibody in a dilution 1/50 (Bäumers et al., 2019). The F-actin microfilaments were stained with phalloidin-TRITC (red), DNA with DAPI (blue). Scale bar in C and D: 200 µm, in F and G: 10 µm.

The BOFs in 5-day-old ds *Chmp4b*-treated females (Figure 2C), were significantly smaller (p < 0.0001; 0.933 ± 0.05 mm; n = 19) compared to dsMock-treated females (1.414 ± 0.03 mm; n = 18) (Figure 2D and E), and showed a fragile appearance. Moreover, ds*Chmp4b*-treated females were not able to ovulate and oviposit. Labeling for CHMP4B in FCs of ds *Chmp4b*-treated females was low (Figure 2F-F’), compared to the labeling in dsMock-treated females (Figure 2G-G’), and it was necessary to overexpose the images in the corresponding channel to detect some labeling in the nucleus of FCs from the treated ovaries (Figure 2F). Labeling of CHMP4B not only appeared in the nuclei of FCs of ds *Chmp4b*-treated insects, it was also found in cytoplasmic vesicles containing condensed DNA (Figure 2F and F’, arrows).

In the BOF of 5-day-old *B. germanica* adults, the follicular epithelium is well-organized, with a uniform distribution, and all the FCs are binucleated since the cytokinesis was arrested (Figure 3A). At this age is when the patency is most apparent (Figure 3B). The contraction of cell membranes had occurred, leaving large intercellular spaces between the FCs (Figure 3B, arrows). During this process, the F-actin microfilaments in the cell membranes display lateral extensions that maintain the contact between cells through microfilament bridges (Figure 3B’, arrowheads). The large endoreplicating nuclei (Figure 3B’’) are located in the center of the cell, occupying almost all the cytoplasm (Figure 3B and B’’).

**Figure 3.**
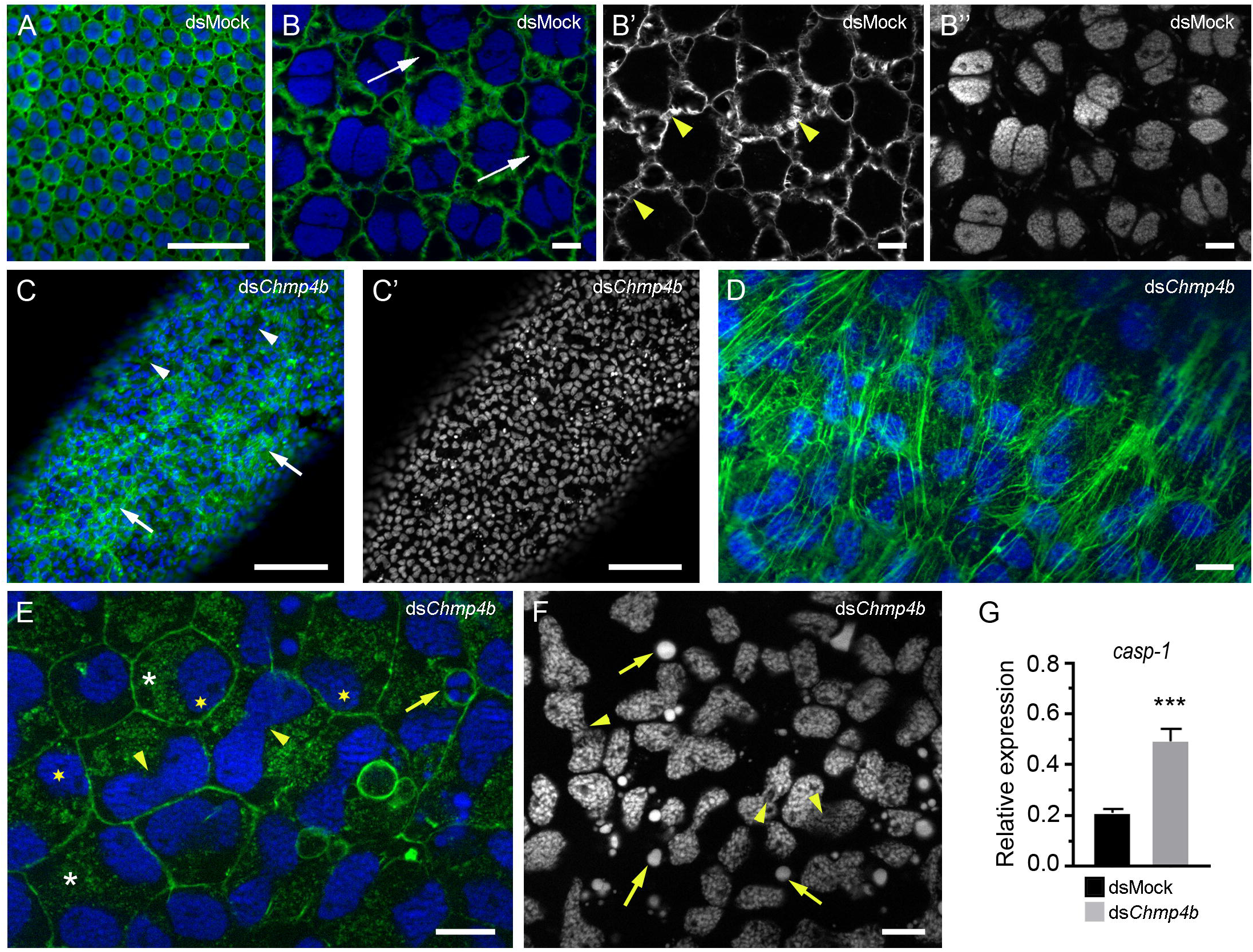
CHMP4B depletion in the basal ovarian follicle of 5-day-old adult *Blattella germanica*. (A) Follicular epithelium from a dsMock-treated female, showing the uniform distribution of the cells. (B) Higher magnification of A, showing the patency and the binucleated FCs. Arrows show the intercellular spaces between the follicular cells. (B’) Actin from B, showing the big intercellular spaces and the bridges of actin connecting the cells (arrowheads). (B’’) Paired nuclei in the FCs. (C) Disorganization of F-actin cytoskeleton in the follicular epithelium ds *Chmp4b*-treated female. The arrows and arrowheads indicate rich and poor actin patches, respectively. (C’) Nuclei from C, showing the variety of size and morphology. (D) Fibers of F-actin in ds *Chmp4b*-treated insects, covering the follicular epithelia. (E) FCs in a ds *Chmp4b*-treated female. The cells are stuck to each other, without signs of patency. Actin staining is abundant in the cytoplasm (asterisks). Vesicles containing rest of DNA are visible (arrow). A high number of cells have nuclei with an hourglass shape (arrowheads). The nuclei appeared stuck to cell membranes (yellow stars). (F) Nuclei from FCs in ds *Chmp4b*-treated females, showing odd shapes. It seems that the nuclei do not complete the karyokinesis and presented bridges connecting both nucleus (arrowheads). Remains of condensate DNA were abundant suggesting apoptosis (arrows). (G) Levels of *casp-1* in ovaries from dsMock- and ds*Chmp4b*-treated females. Data represent copies of *casp-1* mRNA per copy of *eif4aIII* mRNA (relative expression). Data is expressed as the mean ± S.E.M. (n = 6). The asterisks indicate statistically significant differences with respect to dsMock-treated (*** p = 0.0002). The F-actin microfilaments were stained with phalloidin-TRITC (green, except in B’ that appear in white). DNA was stained with DAPI (blue, except in B’’, C’ and F that appear in white). Scale bar in A, C: 100 µm, in B-F: 10 µm.

In 5-day-old ds *Chmp4b*-treated females, the follicular epithelium shows a clear disorganization. The FCs have lost their uniform distribution, showing a remarkable variation of size and shape (Figure 3C and E), affecting the planar polarity of the tissue. Along the oocyte surface, it is possible to visualize areas very rich in F-actin microfilaments (Figure 3C, arrows) beside others with few F-actin fibers (Figure 3C, arrowheads). In those actin-rich areas, the actin microfilaments appear like large bundles of fibers randomly distributed covering the FCs (Figure 3D). Many of the FCs are binucleated, with nuclei of bizarre shapes (Figure 3C’).

The FCs in BOF of ds *Chmp4b*-treated females maintained a narrow contact between them and did not show signs of patency. The F-actin fibers appeared concentrated in the cell membranes without showing expansions (Figure 3E). In addition, it is possible to detect staining for F-actin through the cytoplasm (Figure 3E, asterisks), as well as in the membrane of vesicles containing DNA remains that could suggest the beginning of apoptosis (Figure 3E and F, arrows).

In these 5-day-old *B. germanica* ds*Chmp4b*-treated females, the FCs were smaller, with smaller nuclei than those of dsMock-treated insects and showing odd shapes (Figure 3C’, E, and F). While some cells remained mononucleated (Figure 3E), in others was possible to find two nuclei still attached one to the other with an elongated shape remembering an hourglass (Figure 3E and F, arrowheads), suggesting an incomplete karyokinesis process, as evidenced by visible bridges connecting them. In addition, most of the nuclei lost the central position in the cell and appeared bound to the cytoplasmic membrane (Figure 3E, yellow stars), suggesting changes in the polarity of the cells.

The disorganization of the F-actin cytoskeleton in the FCs of ds *Chmp4b*-treated-adult females, together with the presence of apoptotic vesicles and the changes in the FCs polarity and the morphology of the nuclei, suggested that these cells underwent apoptotic. We measured the expression of *caspase-1* ( *casp-1*) mRNA, a caspase effector, to support this possibility. The *casp-1* expression was significantly increased (131%) in ovaries from ds *Chmp4b*-treated compared to dsMock-treated adults (Figure 3G).

### Expression of genes related to cell planar polarity in ds *Chmp4b*-treated *Blattella germanica* adult ovaries

The morphological changes of FCs in the ds*Chmp4b*-treated females, together with the increase of *actin-5c* expression and F-actin accumulation in the cytoplasm, indicate that the relaxation of the cytoskeleton at the gap junctions that would allow the formation of patency has not occurred. As a result, cells appear tightly connected. All of that drove us to quantify the expression of the genes involved in the cell-cell adhesion regulating the planar polarity in the follicular epithelium.

Based on *D. melanogaster* studies (Casal et al., 2006; Strutt & Strutt, 2021; Thomas & Strutt, 2012), we selected some genes involved in maintaining the epithelial planar polarity and the actin cytoskeleton organization. We measured two cadherins, *ds* and *kug*, and the extracellular serine/threonine protein kinase *fj*, all of them belonging to the *ds*/*fat* system. We also measured the main proteins from the *fz/stan* system, the protocadherin-like wing polarity protein *stan* and *fz*. In addition, we quantified the β-catenin *arm* and the acto-myosin protein *Mhc*.

In ovaries from 5-day-old ds *Chmp4b*-treated females the expression of those genes belonging to *ds*/*fat* system was upregulated (Figure 4A). The expression of *ds* was significantly increased by a 45% (p < 0.05), *kug* expression increased by 99% (p < 0.05) while *fj* increased a 51% its expression, even though not significantly. The genes related with the *fz/stan* system, showed a tendency to increase their expression (Figure 4B). The expression of *stan* increased a 15%, while *fz* increased a 24%. Similarly, the expression of *arm* is significantly upregulated by 21% (p < 0.01, Figure 4C), while *Mhc* tends to be upregulated (29%, Figure 4D).

**Figure 4.**
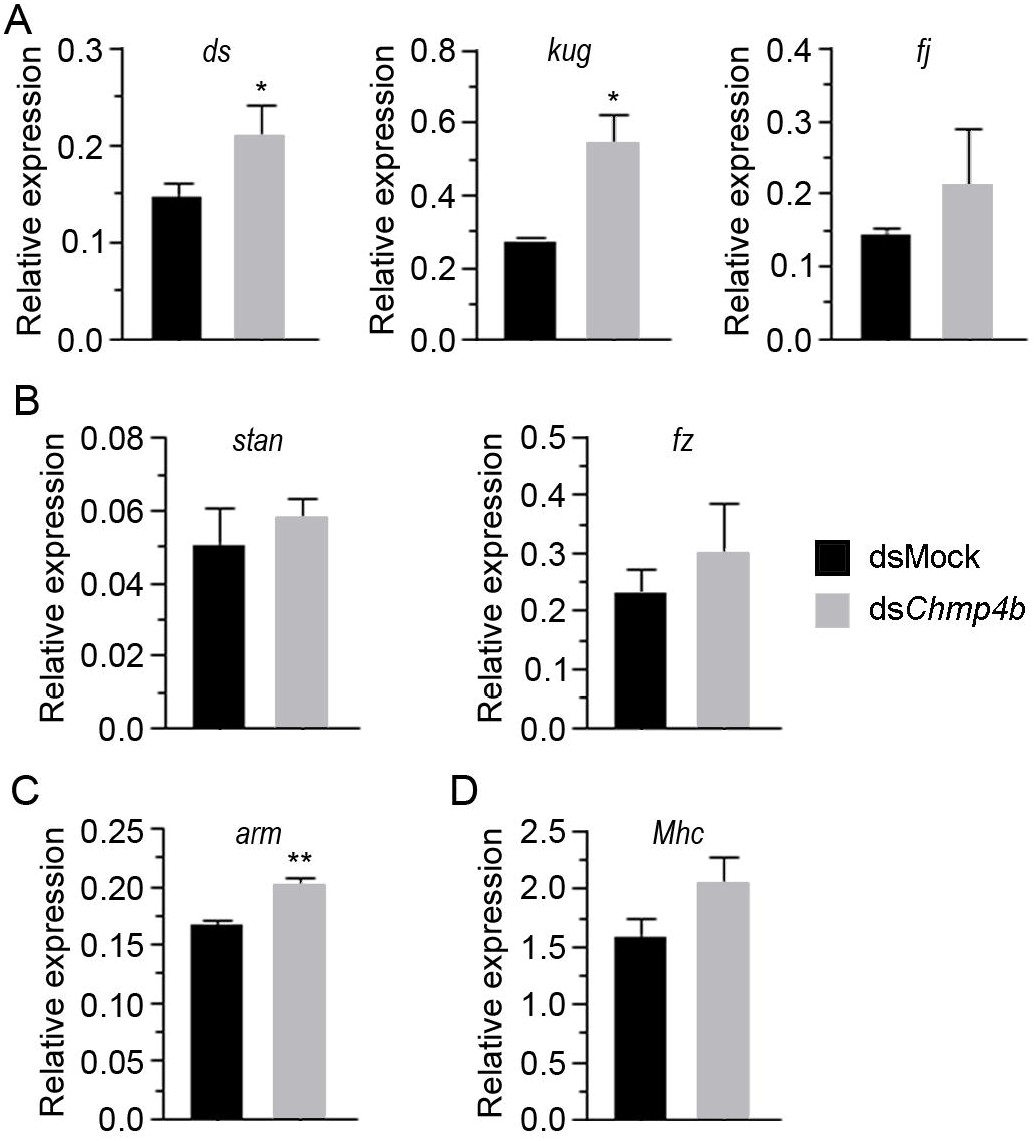
Expression of genes related with planar polarity in ovaries from 5-day-old ds*Chmp4b*-treated adults of *Blattella germanica*. (A) Expression of *dachsous* ( *ds*), *kugelei* (*kug*) and *four-jointed* (*fj*), belonging to *ds*/*fat* system. (B) Expression of *starry night* (*stan*) and *frizzled* ( *fz*), from the *fz*/*stan* system. Expression of (C) the β-catenin *armadillo* (*arm*) and (D) *Myosin heavy chain* (*Mhc*). Data represent copies of mRNA per copy of *eif4aIII* mRNA and are expressed as the mean ± S.E.M. (n = 3). The asterisks indicate statistically significant differences with respect to dsMock: * p = 0.02 ( *ds*); * p = 0.03 (*kug*); ** p = 0.0095 (*arm*).

### Effects of *Chmp4b* depletion on the basal ovarian follicle of *Blattella germanica* last instar nymphs

In the first part of this work, we analyzed the effects of *Chmp4b* depletion in the FCs from ovaries of 5-day-old adult females when cytokinesis is arrested and the change to endoreplication occurs naturally (Irles and Piulachs, 2014). Then, we studied the function of CHMP4B in the ovaries of 6-day-old sixth instar nymphs when the FCs have a high mitotic activity (Irles and Piulachs, 2014).

Newly emerged sixth instar nymphs of *B. germanica* were treated with ds *Chmp4b* or dsMock, and the phenotypes determined were analyzed in nymphal ovaries six days later. The expression levels of *Chmp4b* were significantly depleted (34%, p < 0.01, Figure 5A), while the expression of *actin-5c* tended to be increased (32%; Figure 5B). In these ds*Chmp4b*-treated nymphs, the BOFs were significantly smaller (p < 0.05; 0.25 ± 0.01 mm; n = 5) compared with dsMock-treated nymphs (0.29 ± 0.007 mm; n = 3) (Figure 5C, D, E and F). In some ovarioles from ds *Chmp4b*, the BOFs even had a rounded shape (Figure 5E, S1).

**Figure 5.**
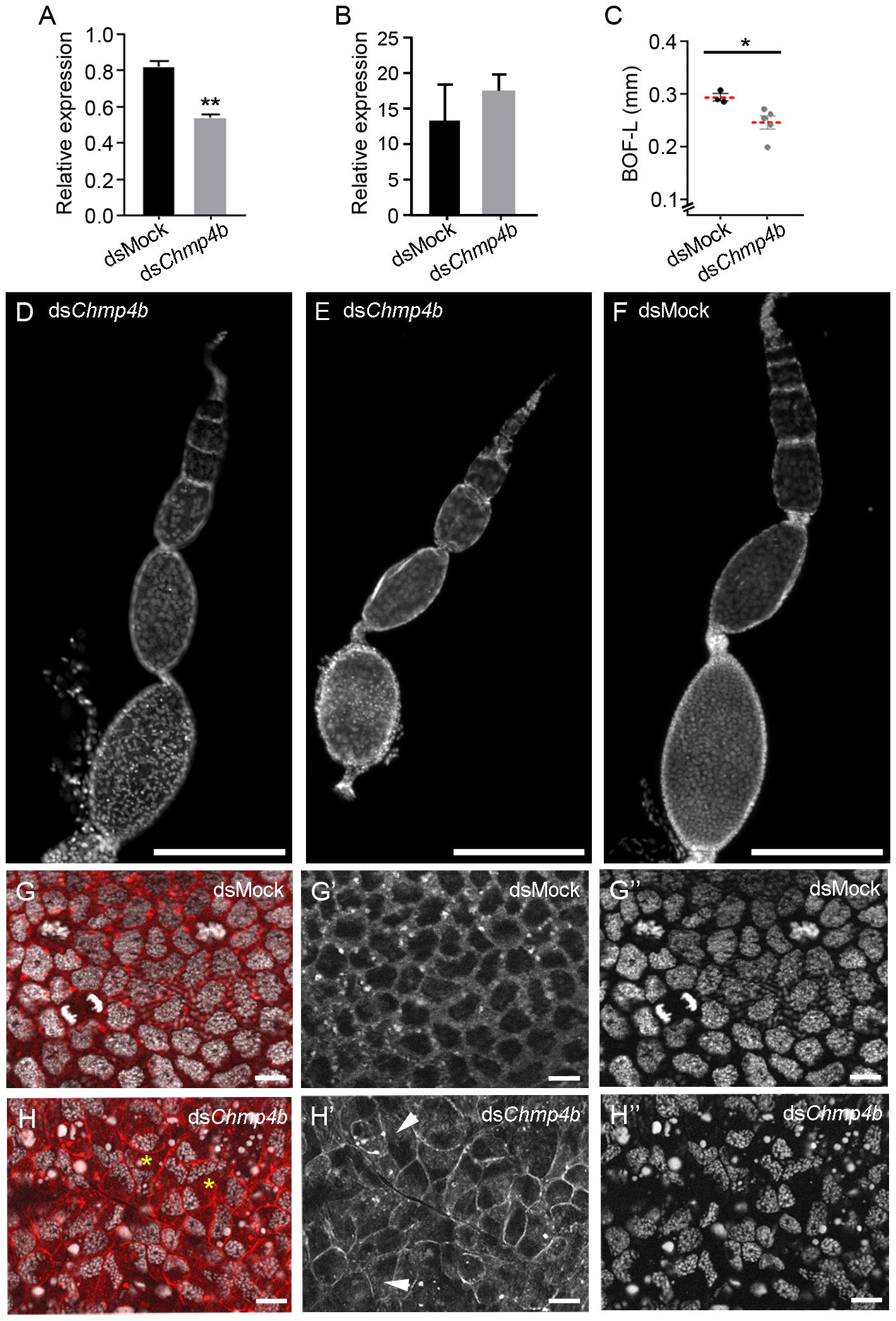
Depletion of CHMP4B in ovaries from 6-day-old sixth-instar nymphs of *Blattella germanica*. Newly emerged last instar nymphs were treated with ds *Chmp4b* or dsMock and analyzed six days later. (A) Relative expression of *Chmp4b*. (B) Relative expression of *actin-5c*. Data represent copies of mRNA per copy of *eif4aIII* mRNA. Data isexpressedasthemean±S.E.M.(n=3).Asterisksindicatestatisticallysignificant differences with respect to dsMock: ** p = 0.0013. (C) Length of BOF (BOF-L; mm) in dsMock- and ds *Chmp4b*-treated nymphs (10 – 20 ovarian follicles were measured per female, n = 3 and 5 females, respectively; * p = 0.03). The dashed line indicates mean. (D) Ovariole from ds *Chmp4b*-treated nymph. (E) Ovariole from ds *Chmp4b*-treated nymph, showing the round shape (see also Figure S1). (F) Ovariole from dsMock-treated nymph. (G) Follicular cells from dsMock-treated nymphs, showing the mitotic figures. (G’) Channel corresponding to F-actin microfilaments of G. (G’’) Channel corresponding to nuclei of FCs of G. (H) Follicular cells of ds *Chmp4b*-treated nymphs. The cell shape and sizes are not uniform, there are not cells under division, and the nuclei of the FCs showed elongated shapes with a polarized position attached to cell membranes (see asterisks). (H’) Channel corresponding to F-actin microfilaments of H, showing the great definition profile of the cell membranes due to the increase F-actin there. The F-actin also appeared distributed by the cytoplasm (arrowheads). (H’’) channel corresponding to nuclei of H, showing the odd nuclei shape. The F-actin microfilaments were stained with phalloidin-TRITC (red in G and H and white in G’ and H’). The DNA was stained with DAPI (white). Scale bar in D-F: 200 µm; G and H: 10 µm.

In the FCs of dsMock-treated *B. germanica* last instar nymphs, the mitotic figures were evident, and all cells were mononucleated, with nuclei of similar size occupying most of the cytoplasm of the cell (Figure 5G and G’’). The F-actin cytoskeleton from the FCs maintains an organized and homogeneous distribution of the cells in a monolayer, without intercellular spaces between them (Figure 5G’). In 6-day-old ds *Chmp4b*-treated nymphs, no mitotic figures were visualized in the follicular epithelium. Most of the FCs were mononucleated as in dsMock, although a few binucleated cells appeared scattered throughout the follicular epithelia (Figure 5H, asterisks). In these ds *Chmp4b*-treated nymphs, the nuclei of FCs had peculiar and elongated shapes (Figure 5H’’), similar to the ones found in ds *Chmp4b*-treated adults (see Figure 3F). The nuclei, in general, are smaller than in dsMock-treated BOFs, and again, they appear polarized on one side of the cells, losing their central position (Figure 5H).

Moreover, in 6-day-old ds *Chmp4b*-treated nymphs, the FCs showed different morphologies and sizes, indicating a loss of planar polarity (Figure S1). The labeling for actin marked sharply the profile of the FCs membranes and filled the cytoplasm not occupied by the nucleus (Figure 5H and H’ arrowheads). It was possible to find DNA vesicles in the cytoplasm of the FCs, indicating cell apoptosis. (Figure 5H and H’’). The increase of F-actin staining is also evident in the oocyte membrane of the BOF (Figure S1) and also in the microtubule organization centers (MTOCs), which usually at this age are sporadically detected in dsMock-treated females, appearing attached to the oocyte membrane in both poles of the oocyte. The MTOCs in BOF of 6-day-old ds *Chmp4b*-treated nymphs do not appear so close to the oocyte membrane, the actin fibers connecting both MTOCs are visible, and some fibers seem to reach the oocyte membrane behind the MTOC (Figure S1).

## DISCUSSION

Insect oogenesis requires a sequential series of changes in the ovary to ensure oocyte maturation and egg formation. To accurately complete all these steps, the organization of cytoskeleton fibers is necessary, and as has been evidenced in *D. melanogaster,* the role of the ESCRT complex in regulating the actin cytoskeleton is essential (Vaccari et al., 2009). Mutations in any component of the ESCRT complex lead to flaws in actin and plasma membrane organization, resulting in multinucleated cells due to cytokinesis failures (Sevrioukov et al., 2005; Vaccari et al., 2009). The planar polarity of these cells was affected, and the disorganization of actin bundles determined spherical egg chambers (Cetera & Horne-Badovinac, 2015). These actin modifications were also described in the testis of ds *Shrb D. melanogaster* -treated adults, where an upregulation of genes related to the cytoskeleton was detected (Chen et al., 2021).

In B. germanica, similarly as it occurs in D. melanogaster (Cetera & Horne-Badovinac, 2015), depletion of Chmp4b provoked an increase in the expression of actin, which resulted in an excess of bundles covering the BOF that adopted, in some cases, a round shape (Figure 5 and S1). In B. germanica, this phenotype was more evident in previtellogenic ovaries from the last instar nymphs than in adults since the ovarian follicles begin to take their shape in the last nymphal instar. Moreover, an accumulation of actin fibers was also evident in the basal oocytes from dsChmp4b-treated nymphs, where both MTOCs and the actin fibers connecting them were frequently visible, while the MTOCs in basal oocytes from control nymphs at the end of the instar, are rarely observed (Figure S1).

In the BOF of *B. germanica* previtellogenic females, levels of *Chmp4b* are higher than in the vitellogenic period. These higher levels of *Chmp4b* coincides with the mitotically active FCs. Conversely, in adults during vitellogenesis, and coinciding with the decrease of *Chmp4b* expression levels, FCs arrest cytokinesis and become binucleated. The depletion of *Chmp4b* mRNA due to dsRNA treatments in both adult and last instar nymphs determines a loss of planar polarity in the ovarian follicular epithelium. The FCs loss their morphology and uniform distribution, appearing closely linked to each other. Furthermore, their nuclei exhibited a loss of polarity, deviating from their central position within the cell and instead becoming attached to the cytoplasmic membrane. The increased expression of proteins belonging *fat/ds* and *fz/stan* systems in adults, related with planar polarity and cell adhesion, would make impossible to relax gap junctions to open spaces between the cells allowing patency to progress. In *D. melanogaster* it was demonstrated that patency requires a local loss of adhesion between FCs for the opening of cell junctions (Isasti-Sanchez et al., 2021). This process relies on the removal of adhesion proteins, including cadherins, from the plasma membrane through endocytic vesicles that requires the presence ESCRT-III complex, and therefore, CHMP4B.

In *B. germanica* last instar nymphs, the FCs in the BOF are mitotically active and mononucleated. Depletion of *Chmp4b* at this stage, leads to incomplete cytokinesis in the FCs, leading to sporadically binucleated cells. Similar to what happens in control adults when *Chmp4b* naturally decreases during the vitellogenesis (Figure 1A). Low levels of *Chmp4b* in adult ovaries are coincident with the cytokinesis arrest in FCs that become binucleated, and undergo endoreplication programming (Irles et al., 2016; Irles & Piulachs, 2014). When depletion of *Chmp4b* is determined early in the adult stage by RNAi treatment, many cells arrest the mitotic program staying mononucleated and not changing their program. In both cases, in nymphs and adults, the FCs can exhibit abnormally elongated nuclei because the karyokinesis has not been completed. Results could be explained by the importance of *Chmp4b* in regulating the timing of membrane abscission (Eikenes et al., 2015; Matias et al., 2015), in a similar way as it was described for vertebrate cells, where depletion of the ESCRT-III components, including *Chmp4b*, produced aberrant nuclei with multiple lobes or micronuclei (Olmos et al., 2015). The consistent results observed in FCs of 6-day-old sixth-instar nymphs and 5-day-old adults of *B. germanica*, after the depletion of *Chmp4b*, characterized by the loss of planar polarity and an incorrect number of nuclei, as well as the presence of aberrant nuclei, suggest that the function of *Chmp4b* is independent of the proliferative stage of the tissue. The changes determined by *Chmp4b* depletion in the FCs affect, in the end, the BOF development, causing female sterility.

In *B. germanica*, CHMP4B is necessary for maintaining the correct rate of FC proliferation, the planar polarity, and the nuclear count of FCs during the BOF growth. CHMP4B is needed to maintain the correct expression of cytoskeleton proteins to allow the BOF to grow and mature properly, incorporating yolk proteins. CHMP4B regulates the correct distribution of actin fibers in the BOF, essential to complete ovulation and oviposition (Alborzi & Piulachs, 2023). All these data help to understand the oogenesis in a basal insect with panoistic ovaries. However, some studies on the other components of the ESCRT-III subcomplex and its regulation will be fundamental to understanding all these processes.

## Abbreviations

BOF: basal ovarian follicle
CHMP4B: charged multivesicular body protein 4B
ESCRT: Endosomal Sorting Complex Required for Transport
FC: follicular cell
MTOC: microtubule organization center
S.E.M.: standard error of the mean

## ACKNOWLEDGMENTS

The authors thank Dr. Thomas Klein, Institut für Genetik, Heinrich-Heine-Universität for kindly supplying a sample of antibody against Shrub. We thank the financial support to the project PID2021-122316OB-I00 from the MCIN/AEI/ 10.13039/501100011033 and by ERDF, a way of making Europe. To the project PID2019-104483GB-I00, funded by MCIN / AEI / 10.13039/501100011033 and to the Catalan Government (2021 SGR 00419). This work was also supported by Pla de Doctorats Industrials de la Secretaria d’Universitats i Recerca del Departament d’Empresa i Coneixement de la Generalitat de Catalunya (grant number 2021 DI 059).

