## Supplementary Table S1 for "CHMP4B contributes to maintaining the follicular cells integrity in the panoistic ovary of the cockroach *Blattella germanica*"

**Table S1.** Primers used for transcript measurements, and for dsRNA synthesis. The length of the amplicon and the accession number of the sequence under study are indicated. F: forward and R: reverse primers

| **Primer set** |  | **primer 5’-3’** | **Amplicon length (bp)** | **Accession number** |
| --- | --- | --- | --- | --- |
| *actin-5c* | F  R | AGCTTCCTGATGGTCAGGTGA  ACCATGTACCCTGGAATTGCCGACA | 213 | AJ862721 |
| *arm* | F  R | CCTGTGTGAACCCTTGGTCT  CACTCTGGAGCCACAACTCA | 100 | PSN33457.1 |
| *casp-1* | F  R | AAGCGGAAGGATTCATACCA  GATGACTGCCTTGCCTCTTC | 80 | LN812812 |
| *Chmp4b* | F  R | ACTACAAAATGAGTTTCTTGGC  GCATTTCCTCAGTTTCTCTTAA | 105 | PSN45592.1 |
| *ds* | F  R | CGGTGAGAATGTTCGTGTGG  CATCGCGTGGGGTATTTTCA | 110 | PSN55018.1 |
| *eIF4aIII* | F  R | ATGGTGACATGCCACAAAAA  GCAACACCTTTCCTTCCAAA | 208 | HF969254 |
| *fj* | F  R | ACAACAACAAGAGGCGTCGT  CTCACAACACTGCCACCTGA | 91 | PSN40108.1 |
| *fz* | F  R | CGCGTGTATGGGTTGGAGTT  ATAGGTCGCTCTGGATATCG | 109 | PSN55438.1 |
| *kug* | F  R | TGCTATGATGCCAAGTCGCT  TGCACCGACTCGTTCACTTT | 90 | PSN45660.1 |
| *Mhc* | F  R | ACACCAGGAAGAACCACCAG  CTGAGTGCCTCAGCCTTACC | 85 | LT717632 |
| *stan* | F  R | ACGCACCGAGATTTTACACC  ATGCTGAAACCAGTGGGAAC | 70 | PSN41253.1 |
| ds*Chmp4b* | F  R | CACAACTGGTGAAGCTATTCAGAA  CTTGTTCCAATTCTTCCAACTCCT | 461 | PSN45592.1 |
| dspolyH | F  R | ATCCTTTCCTGGGACCCGGCA  ATGAAGGCTCGACGATCCTA | 307 | K01149 |
