## Supplementary figures and images for "CHMP4B contributes to maintaining the follicular cells integrity in the panoistic ovary of the cockroach *Blattella germanica*"

### Supplementary Figure S1

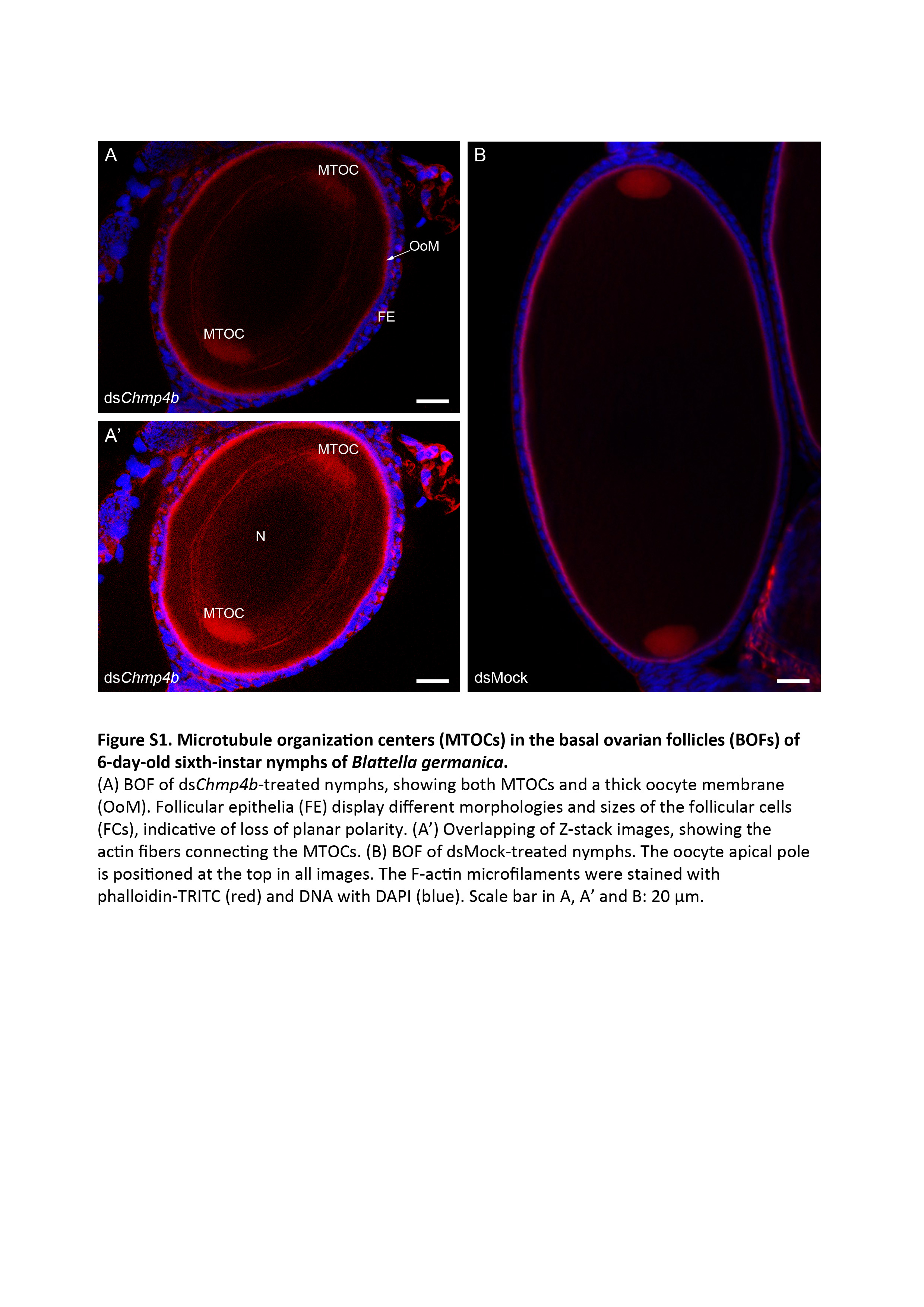
